# Effect of Vinyl Chloride Exposure on Cardiometabolic Toxicity

**DOI:** 10.1101/2021.08.31.458366

**Authors:** Igor N. Zelko, Breandon S. Taylor, Trinath P. Das, Walter H. Watson, Israel D. Sithu, Banrida Wahlang, Marina V. Malovichko, Matthew C. Cave, Sanjay Srivastava

## Abstract

Vinyl chloride is an organochlorine mainly used to manufacture its polymer polyvinyl chloride, which is extensively used in the manufacturing of consumer products. Recent studies suggest that chronic low dose vinyl chloride exposure affects glucose homeostasis in high fat diet-fed mice. Our data suggest that even in the absence of high fat diet, exposure to vinyl chloride (0.8 ppm, 6h/day, 5day/week, for 12 weeks) induces glucose intolerance (1.0 g/kg, i.p) in male C57BL/6 mice. This was accompanied with the depletion of hepatic glutathione and a modest increase in lung interstitial macrophages. Vinyl chloride exposure did not affect the levels of circulating immune cells, endothelial progenitor cells, platelet-immune cell aggregates, and cytokines and chemokines. The acute challenge of vinyl chloride-exposed mice with LPS did not affect lung immune cell composition or plasma IL-6. To examine the effect of vinyl chloride exposure on vascular inflammation and atherosclerosis, LDL receptor-KO mice on C57BL/6 background maintained on western diet were exposed to vinyl chloride for 12 weeks (0.8 ppm, 6h/day, 5day/week). Unlike the WT C57BL/6 mice, vinyl chloride exposure did not affect glucose tolerance in the LDL receptor-KO mice. Plasma cytokines, lesion area in the aortic valve, and markers of lesional inflammation in vinyl chloride-exposed LDL receptor-KO mice were comparable with the air-exposed controls. Collectively, despite impaired glucose tolerance and modest pulmonary inflammation, chronic low dose vinyl chloride exposure does not affect surrogate markers of cardiovascular injury, LPS-induced acute inflammation in C57BL/6 mice, and chronic inflammation and atherosclerosis in the LDL receptor-KO mice.

## Introduction

Environmental pollution accounts for 9 million premature deaths globally, out of which 6 million deaths are attributed to air pollution (1, 2). Studies over the last three decades suggest that air pollution is associated with cardiovascular disease events and cardiovascular mortality, especially myocardial ischemia and stroke (3–5). Population-based studies show that atherosclerosis, the underlying cause of most of myocardial infarction and stroke, is exacerbated by exposure to environmental pollution (4, 5). Studies in experimental animals show that exposure to environmental chemicals such as acrolein augments myocardial ischemic injury (6) and induces dilated cardiomyopathy (7); crotonanaldehyde causes hypotension (8); and acrolein, 1,3-butadiene, and arsenic exacerbate atherosclerosis (9–11). However, little is known about the effect of industrial chemicals on cardiovascular health and injury.

Vinyl chloride (VC) monomer is a colorless and volatile organochlorine mainly used to manufacture the polymer polyvinyl chloride, which is extensively used to make pipes, coating of wire, building materials, and other consumer products. VC is ranked fourth on the Centers for Disease Control and Protection’s Agency for Toxic Substances and Disease Registry Substance Priority List (12). The global demand for VC is more than 18 billion pounds and expected to increase at a rate of 3.5% annually (12). Elevated levels of vinyl chloride are present near the manufacturing facilities, municipal landfills, and at the Environmental Protection Agency Superfund sites, not only as a direct toxin but also as a byproduct of the degradation of other organochlorine compounds such as trichloroethylene (13–16).

VC is a group 1 human carcinogen (17). Major health effects of VC include angiosarcoma of the liver which led to the death of several workers at the manufacturing plant in Kentucky (18). Other adverse health effects of occupational exposure to high levels of VC include arterial hypertension (19), acroosteolysis (20), respiratory irritation with desquamation of the alveolar epithelium (21), ataxia, and dizziness (22), and thrombocytopenia (23, 24). In a cohort analysis of 1658 male workers in Italy, exposure to VC was associated with excessive cardiovascular disease risk (RR=2.25) (25). In our studies, we have found that among highly exposed VC workers the prevalence of steatohepatitis was 80%, indicative of what we call toxicant-associated steatohepatitis, which is characterized by insulin resistance, elevated proinflammatory cytokines but normal liver enzymes (26).

We and others have shown that impaired glucose tolerance and insulin resistance are associated with endothelial dysfunction, endothelial inflammation, and atherosclerosis (27–33). Endothelial dysfunction has also been proposed as a mode of action for the development of hepatic hemangiosarcoma, a sinusoidal endothelial cell tumor that occurs with volatile organic compound exposures including VC (34). Moreover, hepatic steatosis correlates strongly with both hepatic and peripheral insulin resistance (35, 36), and hepatic insulin resistance can lead to an increase in several factors associated with high cardiovascular disease risk including elevations in blood glucose, CRP, small dense LDL, PAI-1, and fibrinogen and low HDL (37). Impaired glucose tolerance is also positively associated with the progression of atherosclerosis and is a prominent risk factor for 5-year changes in carotid atherosclerosis (31, 38) and cardiovascular disease events (39–41). The relative risk for cardiovascular disease events can be increased by up to 40% in subjects with impaired 2h glucose tolerance (31). Pre-clinical studies suggest that diet-induced atherosclerosis is significantly exacerbated in glucose intolerance prone mice as compared with the corresponding glucose intolerance-resistant controls (42). Mechanistic studies suggest that postparandial or transient hyperglycemia induce oxidative stress and subsequently elicit pro-atherogenic responses(43, 44). Thus, VC-induced insulin resistance could potentially induce or exacerbate cardiovascular disease risk.

To minimize the adverse health effects of occupational VC exposure, the Occupational Safety and Health Administration (OSHA) decreased the VC-exposure limit to <1 ppm in 1974 (14, 18). However, very few preclinical studies have been performed to assess the cardiometabolic toxicity of these low-dose VC exposures. Recent studies by our group showed that exposure to VC (0.8 ppm) augments the high fat diet-induced metabolic homeostasis and hepatic toxicity (45, 46). In this study, we examined the plausibility that chronic low dose VC-exposure induces insulin resistance which is sufficient to exacerbate atherosclerosis by promoting endothelial damage, inflammation, and thrombosis.

## MATERIALS AND METHODS

### Animal Exposures

Six-week-old male C57BL/6J mice obtained from Jackson Laboratory (Bar Harbor, ME) were maintained on a low-fat diet (Envigo Teklad Diets, Madison, WI) in a pathogen-free facility accredited by the Association for Assessment and Accreditation of Laboratory Animal Care. All procedures were approved by the University of Louisville Institutional Animal Care and Use Committee. At eight weeks of age, mice were exposed to vinyl chloride at 0.8 ppm or HEPA-filtered air in inhalation chambers for 12 weeks (6 hours per day, 5 days per week) in bedding-free cages as described before (46). Mice were fed a low-fat diet (Envigo Teklad Diets, Madison, WI). To examine the effect of VC on immunotoxicity, mice were intraperitoneally injected with LPS (3 mg/kg) 24 hours before euthanasia. To examine the effect of VC exposure (0.8 ppm, 6 hours/day, 5 days/week for 12 weeks) on atherosclerosis, we used LDL receptor-KO mice on C57BL/6 background and maintained them on a western diet (42% kcal from fat, Harlan Laboratories, Madison, WI). Western diet-fed LDL receptor-KO mice exposed to HEPA-filtered air served as controls. Although LDL receptor-KO mice are mildly hyperlipidemic, a western diet is required to induce atherosclerosis in these mice (47).

### Reagents

Vinyl chloride permeation tubes were obtained from Kin-Tek (La Marque, TX). Primers and probes for real-time PCR were purchased from Integrated DNA Technologies (Coralville, IA) and ThermoFisher Scientific (Waltham, MA). Sources of antibodies used for western blotting were: anti-catalase, -SOD2, -HO1, -cleaved caspase 1 p20, -NRF2 (Santa Cruz Biotechnology, Dallas, TX); anti-SOD1 (GeneTex, Irvine, CA); anti-SOD3 (Novus, Centennial, CO). Sources of antibodies for the flow cytometry include: FITC-anti-Sca-1 (Ly-6A/E), APC-anti-Flk1 (CD309), APC-eFluor780-anti-CD41, PE-Cyanine7-anti-Sca-1, FITC-anti-Nk1.1, PE-anti-Ly6C, PerCPe710-anti-CD8, PECy7-anti-CD62, APC-anti-CD19, Alexa700-anti-Gr-1, APCe780-anti-CD3, eVolve605-CD11b, and e650-anti-CD4 antibodies were purchased from eBioscience (San Diego, CA). Lipopolysaccharides from Escherichia coli O111:B4 (LPS) were obtained from Sigma-Aldrich Cat# 2630. Fc Block (CD32/CD16) (Leinco Technologies; St. Louis, MO), counting beads (Spherotech; Lake Forest, IL), BSA (Rockland Immunochemicals, Limerick, PA). All other chemicals and enzymes were from Sigma Chemical Co. (St. Louis, MO), or Invitrogen (Carlsbad, CA).

### Glucose Tolerance Test

Glucose tolerance tests were performed after a 6-hour fast by injecting D-glucose (1 g/kg; i.p.) in sterile saline as described (48, 49).

### Complete Blood Counts

Complete blood counts (CBCs) were measured on Hemavet 950FS hematology analyzer as described (48, 49).

### Cytokine Analyses

Plasma IL-6 levels were measured by mouse IL-6 ELISA kit (ThermoFisher Scientific, Waltham, MA).

### Flow Cytometric Analyses of Circulating Endothelial Progenitor Cells, Immune Cells, and Platelet-Leukocyte Adducts

Blood endothelial progenitor cells (EPCs; Flk-1+/Sca-1+ cells), immune cells, and platelet-leukocyte aggregates were analyzed on a BD-LSRII flow cytometer as described (48, 49).

### Atherosclerotic lesion analysis

Atherosclerotic lesions in the aortic valves were quantitated as described before (9, 33, 50–55). Adhesion of immune cells to mouse aortic endothelial cells (MAEC) *in vitro* was performed in 2-chloroacetaldehyde (CA) and 2-chloroethanol (CE) as described before (56).

### Bronchoalveolar Fluid and Lung Cell Isolation

Bronchoalveolar Fluid (BALF) was isolated as described previously (57). Briefly, mice were anesthetized using pentobarbital and a catheter connected to 1 mL syringe was inserted into the trachea. Ice-cold PBS with 0.1 mM EDTA was administered into the lungs and then removed slowly. This procedure was repeated two more times to achieve maximum recovery of lung macrophages. Cells were pelleted by centrifugation, resuspended in PBS/BSA, and used for flow cytometry analysis.

Lung single-cell suspensions were prepared using Lung Dissociation Kit (Miltenyi, Auburn, CA) according to manufactures protocol. Briefly, lungs were perfused with saline via the right ventricle. The lungs were excised and the left lobe was minced with scissors, transferred into C-tubes (Miltenyi, Auburn, CA), and processed in digestion buffer in a GentleMACS dissociator (Miltenyi, Auburn, CA) according to the manufacturer’s instructions. Digested lungs were passed through a 40-μm nylon mesh strainer to obtain a single-cell suspension. Cells were counted using a TC20 Automated Cell Counter (BioRad, Hercules, CA).

### Quantitative RT-PCR and Western Blotting

Total RNA was prepared from frozen lung and liver tissues using Quick-RNA MiniPrep Kit (Zymo Research, Irvine, CA), and quantitative RT-PCR was performed as described (58). For western blotting, tissue homogenates were prepared in RIPA buffer (50 mM Tris-HCl pH8.0, 0.15 M NaCl, 0.1% SDS, 0.5% sodium deoxycholate, 1% NP-40) with protease and phosphatase inhibitors, and the proteins were separated on a 8-16% SDS-PAGE gel (Invitrogen, Carlsbad, CA) and probed with appropriate antibodies (59).

### Glutathione Levels

Mouse liver and lung tissues were pulverized in liquid nitrogen and transferred to a centrifuge tube containing an equal volume of ice-cold 10%□(w/v) perchloric acid, 0.2□M boric acid, and 20□μM γ-glutamyl glutamate as an internal standard (60). Extracts were centrifuged at 16,000□×g for 2 minutes to remove the precipitated protein. The protein-free extracts were derivatized with iodoacetic acid and dansyl chloride and analyzed by HPLC (Waters Corporation, Millford, MA) as previously described (61). Concentrations of thiols and disulfides were determined by integration relative to the internal standard.

### Malonaldialdehyde Quantitation

The levels of malondialdehyde (MDA) in the liver and lung were measured as described previously (62). Briefly, 20 mg tissue was homogenized in 400 μL water containing a final concentration of 400 μM EDTA, 20 μM butylated hydroxytoluene (BHT), and 20 μM desferal to inhibit the formation of aldehydes during sample homogenization and processing. Samples were derivatized with 2, 3, 4, 5-pentafluorobenzyle bromide for 1h at 80 °C, extracted with hexane, and analyzed using gas chromatography-mass spectrometry as described (63).

### Statistical Analyses

All values are presented as means ± SEM of data from *n* independent experiments, as indicated in the figure legends. The statistical significance of differences was determined by t-test. A one-way analysis of variance (ANOVA) with Holm-Sidak *post hoc* test was used to compare differences between multiple treatment groups. P-value of <0.05 indicated statistically significant differences. All analyses were performed using Excel and GraphPad Prism software (GraphPad Software, San Diego, CA).

## RESULTS

### Vinyl Chloride Induces Glucose Intolerance

Recent studies show that short term (6 weeks) VC exposure moderately increases oral glucose intolerance in low fat diet-fed mice and appreciably exacerbates western diet-induced glucose intolerance (46). However, the effect of long-term VC exposure (12 weeks) on glucose homeostasis, especially in the absence of western diet remains unclear. We observed that in the low fat-diet-exposed mice VC did not affect the weight gain (**Fig. 1A**), blood glucose (**Fig. 1B**), insulin levels (**Fig. 1C**), or hepatic expression of genes associated with glucose homeostasis (**Fig. 1D**; glutamine synthetase, glucokinase, glucose-6-phosphatase, phosphoenolpyruvate 1, glycogen synthase kinase 3α, glucose transporter-2, and glucose transporter-3, and insulin-like growth factor-1). However, when subjected to a glucose tolerance test (GTT), the VC-exposed mice showed a slower glucose clearance rate as compared with the air-exposed controls (**Fig. 1 E-F**). These data suggest that chronic VC exposure augments glucose intolerance in low fat diet-fed mice.

**Figure 1:**
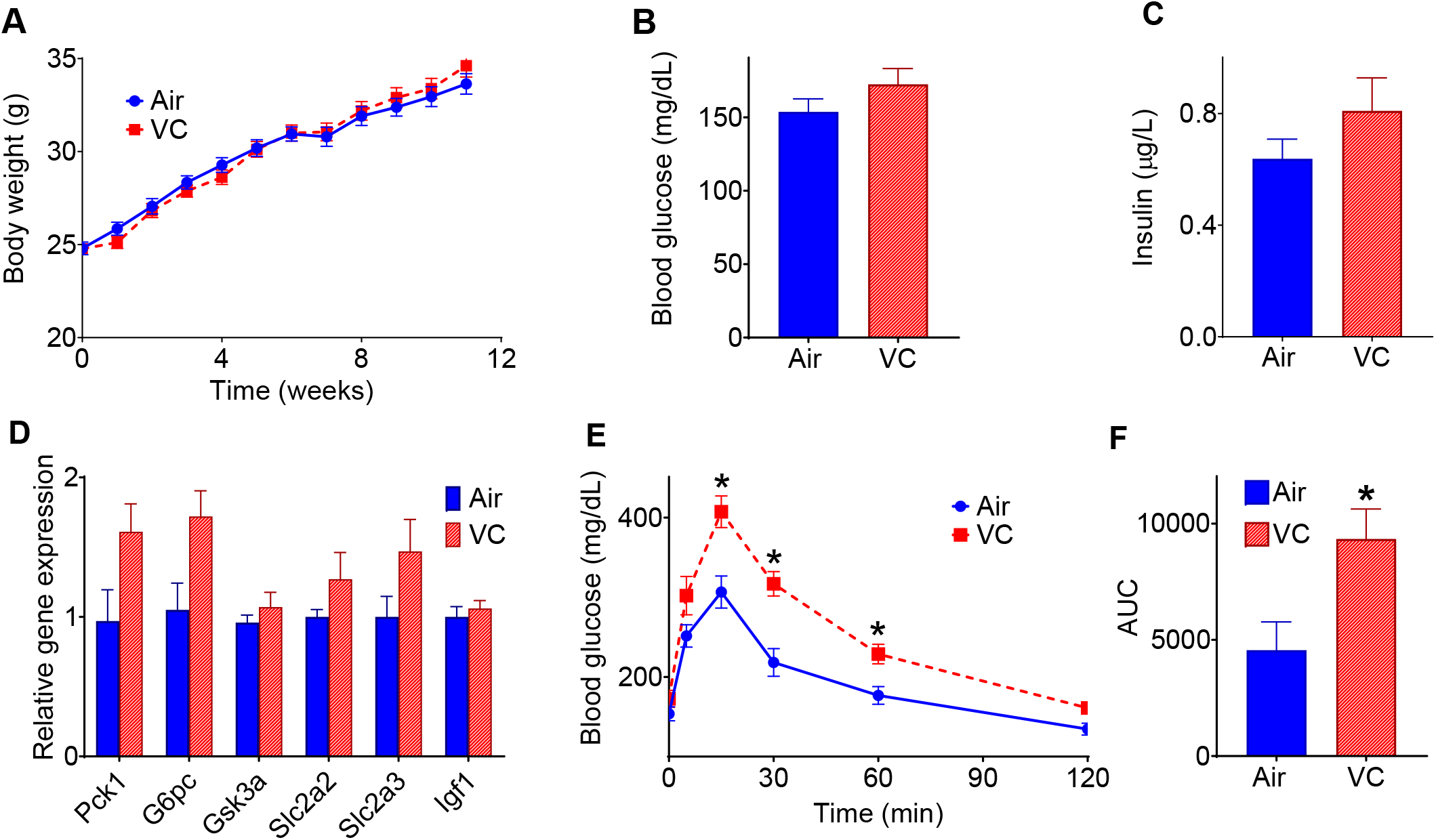
Metabolic changes in vinyl chloride-exposed mice. (**A**) Mice were exposed to HEPA-filtered air (Air) or vinyl chloride (0.8 ppm) for 12 weeks, and body weights were measured at indicated time points (*n* = 25/group). (**B**) Blood glucose level at euthanasia. (**C**) Plasma insulin level at euthanasia. (**D**) The hepatic expression of genes involved in maintaining glucose homeostasis. (**E**) Glucose (1 g/kg; i.p.) tolerance test (GTT) in mice exposed to VC for 12 weeks. (**F**) The area under the curve (AUC) for GTT (*n* = 8/group). Values are mean ± SEM. \**P* < 0.05 *vs* air-exposed controls.

### Hepatic Oxidative Stress and Inflammation in Vinyl Chloride-Exposed Mice

We have recently shown that benzene, another volatile organic compound, induces systemic insulin resistance in a hepatic oxidative stress-dependent manner (64). Similar to benzene exposure, we observed that chronic VC exposure depleted the total and reduced hepatic glutathione levels (**Fig. 2A**), but did not affect the levels of cysteine (Cys), cystine (CySS), oxidized glutathione (GSSG), and GSH/GSSG ratio (**Fig. 2A**). To assess whether depletion in hepatic glutathione is driven by oxidative stress or conjugation with VC metabolites for excretion, we measured the levels of lipid peroxidation-derived aldehyde, malonaldialdehyde (MDA), a hallmark of oxidative stress. As shown in **Fig. 2B**, the hepatic MDA levels in VC-exposed mice were also comparable to controls, suggesting that VC-induced depletion in hepatic glutathione levels is likely a reflection of conjugation of VC metabolites with glutathione rather than overt oxidative stress. Measurement of the abundance of oxidative defense enzymes in the liver showed that VC exposure significantly depletes hepatic superoxide dismutase 3 (SOD3) and heme oxygenase 1 (HO1) levels but does not affect the levels of SOD1, SOD2, catalase (CAT), and nuclear factor erythroid-derived 2 (NRF2) (**Fig. 2C**). Together, these data suggest that although VC exposure does not induce overt oxidative stress, some pathways of the oxidative defense system are compromised.

**Figure 2:**
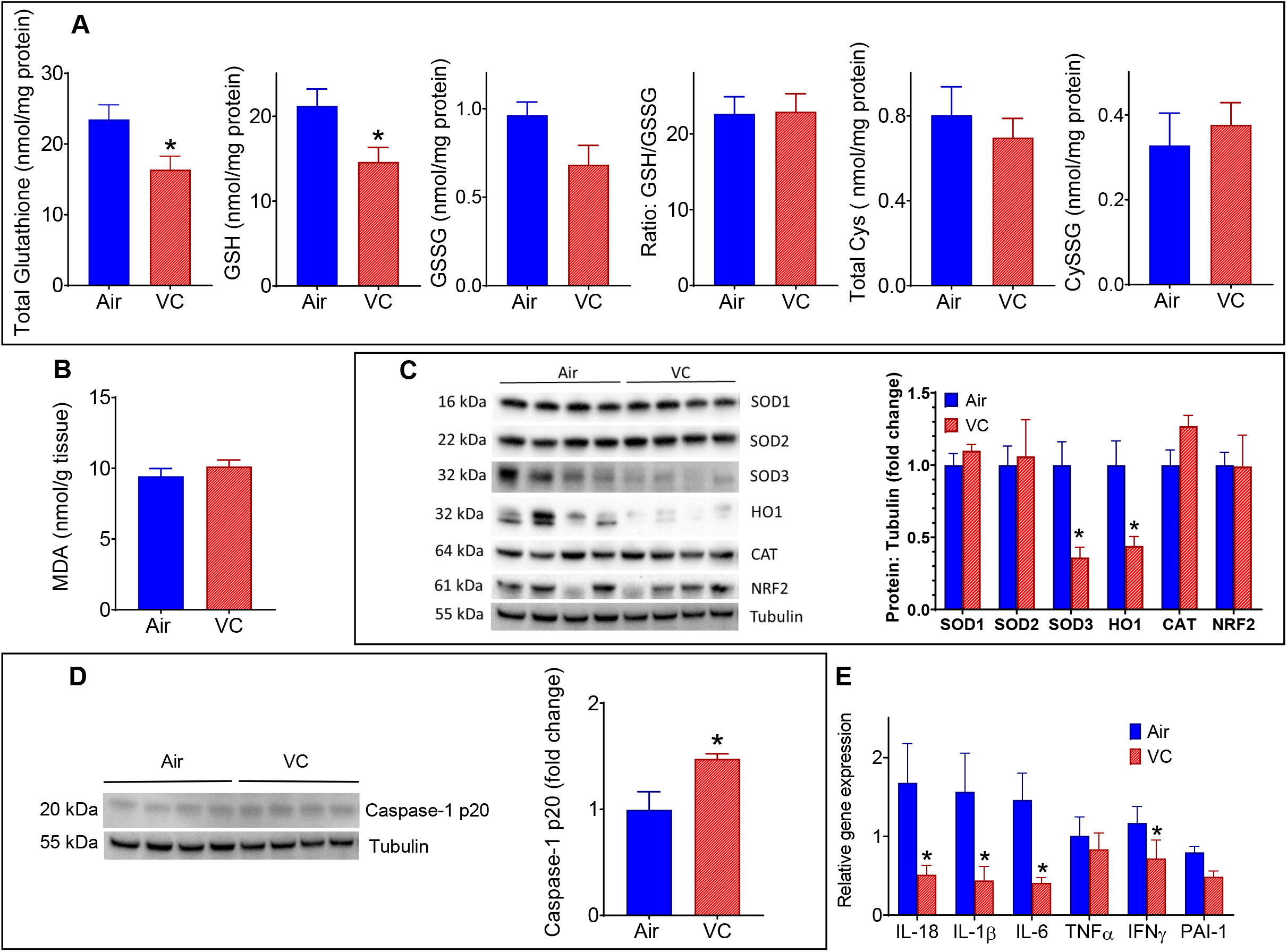
Hepatic oxidative stress and inflammation in vinyl chloride-exposed mice. **(A)** Total glutathione, reduced glutathione (GSH), oxidized glutathione (GSSG), cysteine (Cys) and cystine (CySS) levels in the liver of VC-exposed male C57BL/6 mice. (**B**) Hepatic malondialdehyde (MDA) levels. (**C**) Levels of oxidative defense and antioxidant proteins in the liver. (**D**) Abundance of Caspase-1 in the liver. (**E**) Hepatic expression of cytokines. Values are mean ± SEM. \**P* < 0.05 *vs* air-exposed controls.

Toxicant-induced imbalance in the hepatic oxidative defense system can induce hepatic inflammation, which in turn, can trigger insulin resistance (64). Our data show that VC-exposure induced the hepatic levels of Caspase-1 protein (**Fig. 2D**), the precursor of the inflammatory cytokines IL-1β and IL-18. However, paradoxically the mRNA levels of IL-1β, IL-18 and some of the other pro-inflammatory cytokines (**Fig. 2E**) were significantly lower in the VC-exposed mice. The underlying cause(s) of this discrepancy is unclear.

Pulmonary Inflammation and Oxidative Stress in Vinyl Chloride-Exposed Mice: Since the lung is the first target of inhaled toxicants, we examined the effect of inhaled VC on pulmonary inflammation and oxidative stress. Measurement of tissue resident immune cells in the lungs of VC-exposed mice by flow cytometry (**Fig. 3A**) showed a two-fold increase in the levels of interstitial macrophages as compared with the air-exposed mice (**Fig. 3B**). VC exposure did not affect the abundance of alveolar macrophages, eosinophils, neutrophils, CD3^+^ T-cells, and CD19^+^ B-cells (**Fig. 3 Bi** and **Fig. 3 Bii**). In the bronchoalveolar lavage fluid (BALF), VC exposure neither affected the levels of interstitial and alveolar macrophages nor the polarization of alveolar macrophages (**Fig. 3C** and **Fig 3D**). Indices of pulmonary oxidative stress such as total and reduced glutathione, MDA, and oxidative defense enzymes in the VC-exposed mice were comparable with the air-exposed controls (**Fig. 3E**).

**Figure 3:**
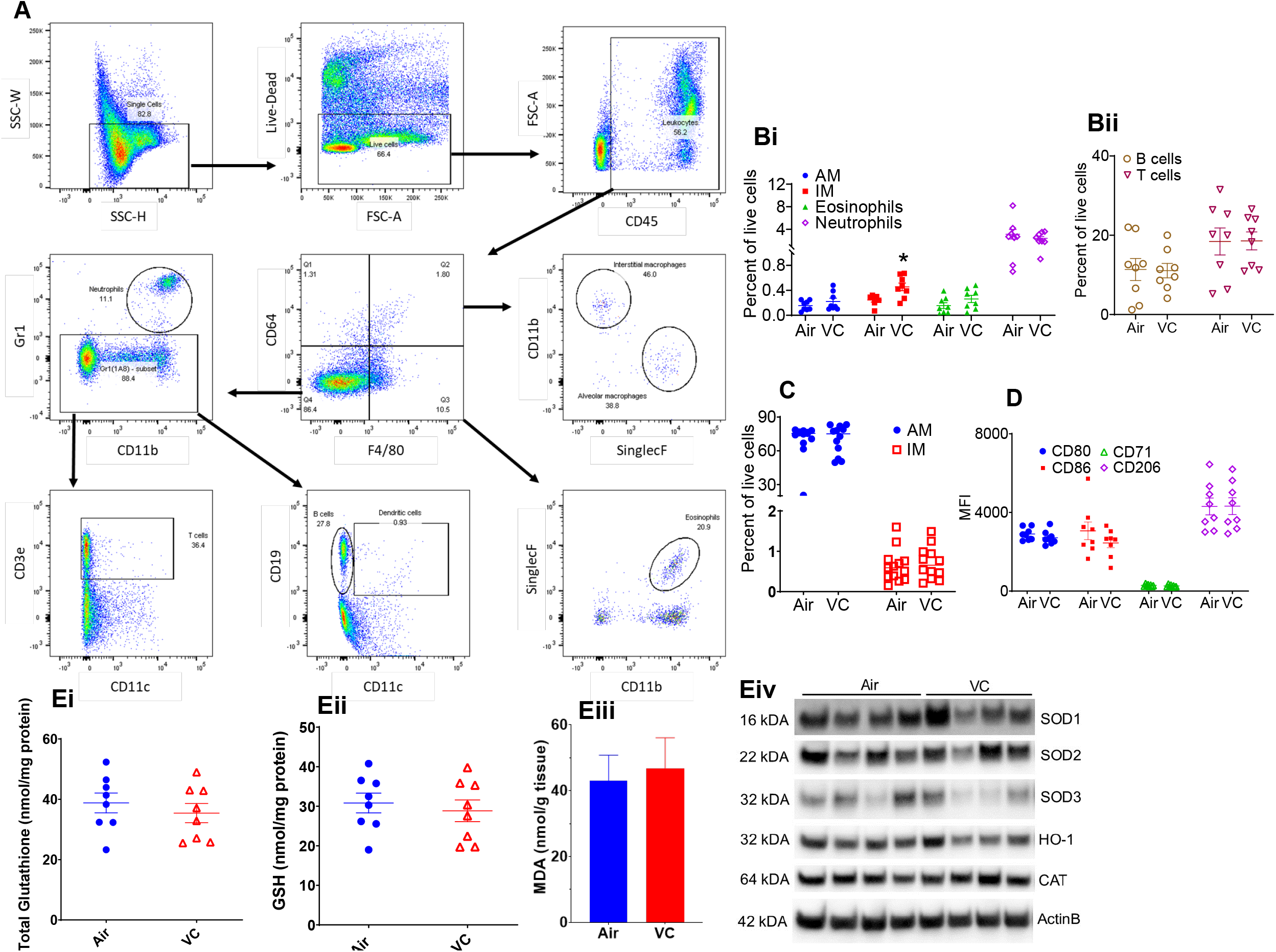
Pulmonary inflammation in vinyl chloride-exposed mice. (**A)** Gating scheme for the flow cytometric analysis of immune cells in the lungs and BALF of VC-exposed male C57BL/6 mice. (**B**) Levels of immune cells – alveolar macrophages, AM; Interstitial macrophages IM; Eosinophils; and Neutrophils (**Bi**) and B-cells and T-cells (**Bii**) in the lungs. (**C**) Abundance of macrophages in the bronchoalveolar lavage fluid (BALF) of VC-exposed male C57BL/6 mice. (**D**) Analysis of macrophage polarization markers in BALF. (**E**) Indices of pulmonary oxidative stress: Total glutathione (**Ei**), GSH (**Eii**), MDA (**Eiii**) and western blots of oxidative defense enzymes (**Eiv**). Values are mean ± SEM. \**P* < 0.05 *vs* air-exposed controls.

### Effect of Chronic Vinyl Chloride Exposure on the Surrogate Markers of Cardiovascular Toxicity

Hyperglycemia or insulin resistance can affect endothelial functions. Therefore, to assess the effect of chronic VC exposure on vascular toxicity, we measured the levels of endothelial progenitor cells (EPCs) in the peripheral blood. Depletion in circulating EPCs is associated with increased cardiovascular disease (65) and could foretell potential future cardiovascular events (66). In this study, we observed that although 12 weeks of exposure to VC significantly decreases the circulating levels of progenitor cells (Sca^+^), it did not affect the blood EPCs (Flk^+^/Sca^+^; **Table 1**). Levels of EPCs in the bone marrow of VC-exposed mice were also comparable with air-exposed controls (data not shown).

**Table 1:** Parameters Measured in the Blood.

| Parameters | Air | VC | Air + LPS | VC + LPS |
| --- | --- | --- | --- | --- |
| **Complete Blood Counts** |  |  |  |  |
| WBC/ $\mu\text{L}$ | 1783 ± 337 | 1428 ± 155 | 1654 ± 152 | 1514 ± 96 |
| Neutrophils/ $\mu\text{L}$ | 805 ± 341 | 368 ± 118 | 1017 ± 127 | 1017 ± 78 |
| Lymphocytes/ $\mu\text{L}$ | 928 ± 110 | 1031 ± 96 | 517 ± 65*<sup>§</sup> | 399 ± 28*<sup>§</sup> |
| Monocytes/ $\mu\text{L}$ | 43 ± 10 | 27 ± 6 | 89 ± 26<sup>§</sup> | 66 ± 7 |
| Eosinophils/ $\mu\text{L}$ | Traces | Traces | Traces | Traces |
| Basophils/ $\mu\text{L}$ | Traces | Traces | Traces | Traces |
| Platelets ($10^3/\mu\text{L}$) | 825 ± 70 | 757 ± 23 | 396 ± 36*<sup>§</sup> | 375 ± 27*<sup>§</sup> |
| Mean platelet volume (fL) | 4.48 ± 0.14 | 4.31 ± 0.04 | 4.99 ± 0.1*<sup>§</sup> | 4.91 ± 0.08*<sup>§</sup> |
| RBC ($10^6/\mu\text{L}$) | 9.4 ± 0.2 | 9.2 ± 0.1 | 9.6 ± 0.3 | 9.5 ± 0.3 |
| Hemoglobin (g/dL) | 12.4 ± 0.1 | 12.6 ± 0.2 | 13.2 ± 0.4 | 13.1 ± 0.4 |
| Hematocrit (%) | 40.1 ± 0.4 | 40.3 ± 0.7 | 41.5 ± 1.4 | 41.3 ± 1.4 |
| Mean corpuscular volume (fl) | 43.2 ± 0.4 | 43.9 ± 0.4 | 43.1 ± 0.5 | 43.2 ± 0.3 |
| Mean corpuscular hemoglobin (pg) | 13.4 ± 0.1 | 13.6 ± 0.1 | 13.7 ± 0.1 | 13.8 ± 0.1 |
| Mean corpuscular hemoglobin concentration (g/dL) | 30.9 ± 0.2 | 31.3 ± 0.4 | 31.8 ± 0.3 | 31.8 ± 0.3 |
| Red cell distribution (%) | 18.3 ± 0.3 | 18 ± 0.2 | 18.4 ± 0.3 | 17.9 ± 0.1 |
| **Circulating Immune Cells (Measured by Flow Cytometry)** |  |  |  |  |
| NK1.1<sup>+</sup> Cells/$\mu\text{L}$ | 15 ± 2 | 16 ± 4 | 6 ± 1<sup>§</sup> | 5 ± 1*<sup>§</sup> |
| CD19<sup>+</sup> B-Cells/$\mu\text{L}$ | 148 ± 28 | 143 ± 21 | 39 ± 6*<sup>§</sup> | 38 ± 6*<sup>§</sup> |
| CD4<sup>+</sup> T-Cells/$\mu\text{L}$ | 46 ± 8 | 48 ± 5 | 13 ± 2*<sup>§</sup> | 16 ± 1*<sup>§</sup> |
| CD8<sup>+</sup> T-Cells/$\mu\text{L}$ | 0.22 ± 0.03 | 0.23 ± 0.02 | 0.52 ± 0.12*<sup>§</sup> | 0.4 ± 0.07 |
| GR1<sup>+</sup> Granulocytes/$\mu\text{L}$ | 69 ± 12 | 69 ± 6 | 558 ± 86*<sup>§</sup> | 407 ± 38*<sup>§</sup> |
| **Circulating Progenitor Cells (Measured by Flow Cytometry)** |  |  |  |  |
| Sca<sup>+</sup> per 1000 small cells | 4.4±0.6 | 2.9±0.3* | ND | ND |
| Flk<sup>+</sup>/Sca<sup>+</sup> per 1000 small cells | 0.8±0.3 | 0.4±0.1 | ND | ND |
| **Platelet Activation Markers (Measured by Flow Cytometry)** |  |  |  |  |
| Leukocyte- platelet aggregates (% of leukocytes) | 41.8 ± 1.8 | 38.2 ± 1.4 | ND | ND |
| Monocytes- platelet aggregates (% of monocytes) | 40.8 ± 2.0 | 37.8 ± 1.4 | ND | ND |
| Lymphocytes- platelet aggregates (% of lymphocytes) | 30.3 ± 0.7 | 29.6 ± 0.8 | ND | ND |

VC-Vinyl chloride. N=8-13 per group. Values are mean ± SEM. \*P<0.05 vs Air. <sup>§</sup>p<0.05 vs VC

Compromised glucose homeostasis can also affect thrombosis (67, 68). Therefore, to examine the effect of chronic VC exposure on markers of thrombosis, we measured platelet activation, the key event in the etiology of thrombosis. Our data show that markers of platelet activation as assessed by the circulating levels of the platelet aggregation with leukocytes, monocytes, or lymphocytes in the peripheral blood of VC-exposed mice were comparable with the air-exposed controls (**Table 1**).

Perturbation in glucose handling could also induce systemic inflammation, and pro-inflammatory cytokines are prognosticators of future cardiovascular events (69). However, little is known about the effect of chronic VC exposure on immune functions. We observed that 12 weeks of exposure to VC did not affect CBC (**Table 1**). Flow cytometric analyses of circulating immune cells showed that levels of NK1.1+ natural killer cells, CD19^+^ B-cells, CD4^+^ T-cells, CD8^+^ T-cells, and Gr1^+^ granulocytes in VC-exposed mice were comparable to air-exposed controls (**Table 1**). VC exposure also did not affect plasma levels of pro-inflammatory cytokines such as IL-6 (**Fig. 4A**). Collectively, our data suggest that VC-induced moderate glucose intolerance did not affect the surrogate markers of cardiovascular injury and toxicity.

**Figure 4:**
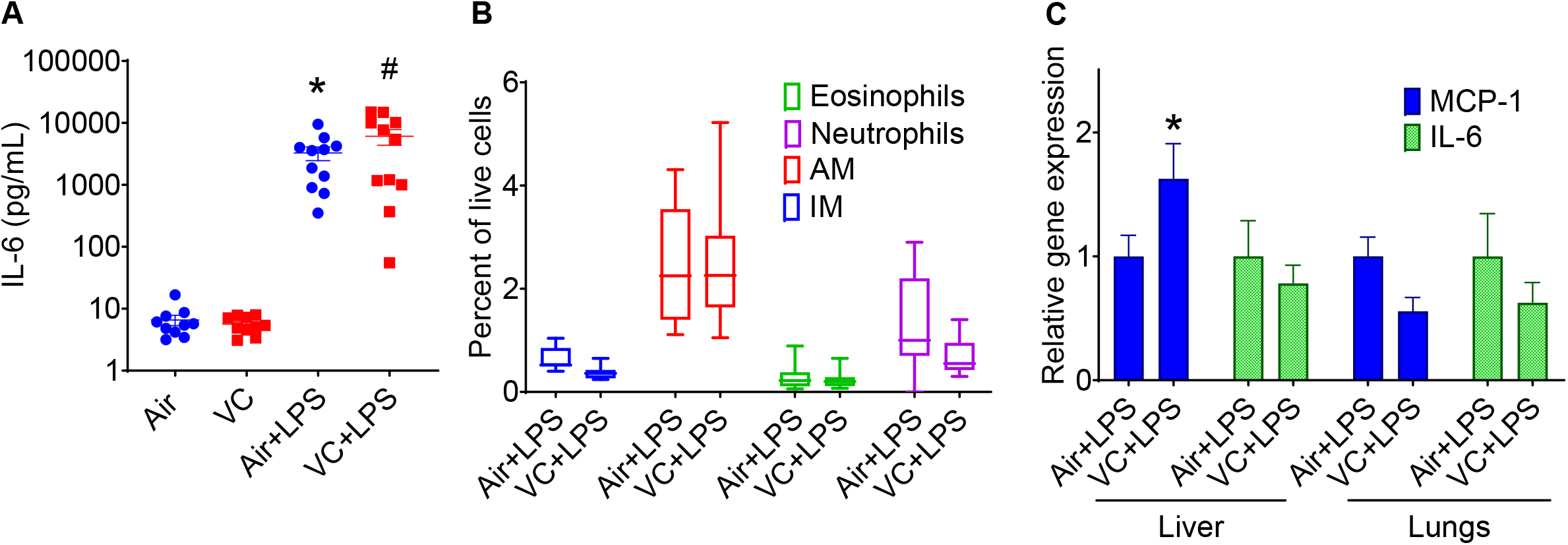
Effect of lipopolysaccharide on pulmonary inflammation in vinyl chloride-exposed mice. Mice exposed to VC or filtered air were treated with LPS (3mg/kg) 24h before euthanasia. (**A)** Plasma IL-6 levels. (**B**) Levels of immune cells in the lungs of LPS-treated mice. (**C**) Expression of IL-6 and MCP-1 in the lungs and liver of LPS-treated mice. Values are mean ± SEM. \**P* < 0.05 *vs* air-exposed controls.

### Sensitivity of Vinyl Chloride-exposed Mice to Acute Inflammation

To examine whether VC exposure affects the sensitivity to an acute secondary hit by an inflammatory stimulus, we exposed a sub-set of VC- and air-exposed mice to LPS (24h before euthanasia) and measured the markers of inflammation in the liver, the lung, and the plasma. Our data show that, as expected, LPS augmented the circulating levels of cytokines such as IL-6 (**Fig. 4A**), but the plasma levels of IL-6 in VC-LPS-exposed mice were comparable with air-LPS-exposed mice. LPS did not affect the levels of inflammatory cells in the lungs (**Fig. 4B**), and cytokine (IL-6) and chemokine (MCP-1) levels in the lungs, but increased the abundance of MCP-1 by 1.7-fold in the liver (**Fig. 4C**). Together, these data suggest that LPS significantly increases the levels of MCP-1 in the liver of VC-exposed mice.

### Sensitivity of Vinyl Chloride-exposed Mice to Chronic Inflammation

Next, we examined whether VC exposure renders the mice more sensitive to chronic and sustained inflammation such as atherosclerosis. Previous studies have shown that exposure to environmental pollutants such as particulate matter, acrolein, and arsenic exacerbates atherosclerosis in hyperlipidemic mice (9, 11, 70). Moreover, insulin resistance aggravates atherosclerosis (71, 72), and deficiency of MCP-1 prevents atherosclerosis in experimental animals (73, 74). Our *in vitro* studies show that incubation of mouse aortic endothelial cells with VC metabolites 2-chloroacetaldehyde and 2-chloroethanol enhance the adhesion of immune cells to the endothelial cells (**Fig. 5A**). However, 12 weeks of inhalation of VC to pro-atherogenic LDL receptor-knockout male mice did not affect the body weight (**Fig. 5B**), glucose tolerance (**Fig. 5C-E**), or plasma cholesterol (**Fig, 5F**), and atherosclerotic lesion area in the aortic valves (**Fig. 5G**) and the innominate artery. These data suggest that chronic inflammatory conditions such as atherosclerosis do not affect the vascular toxicity of VC.

**Figure 5:**
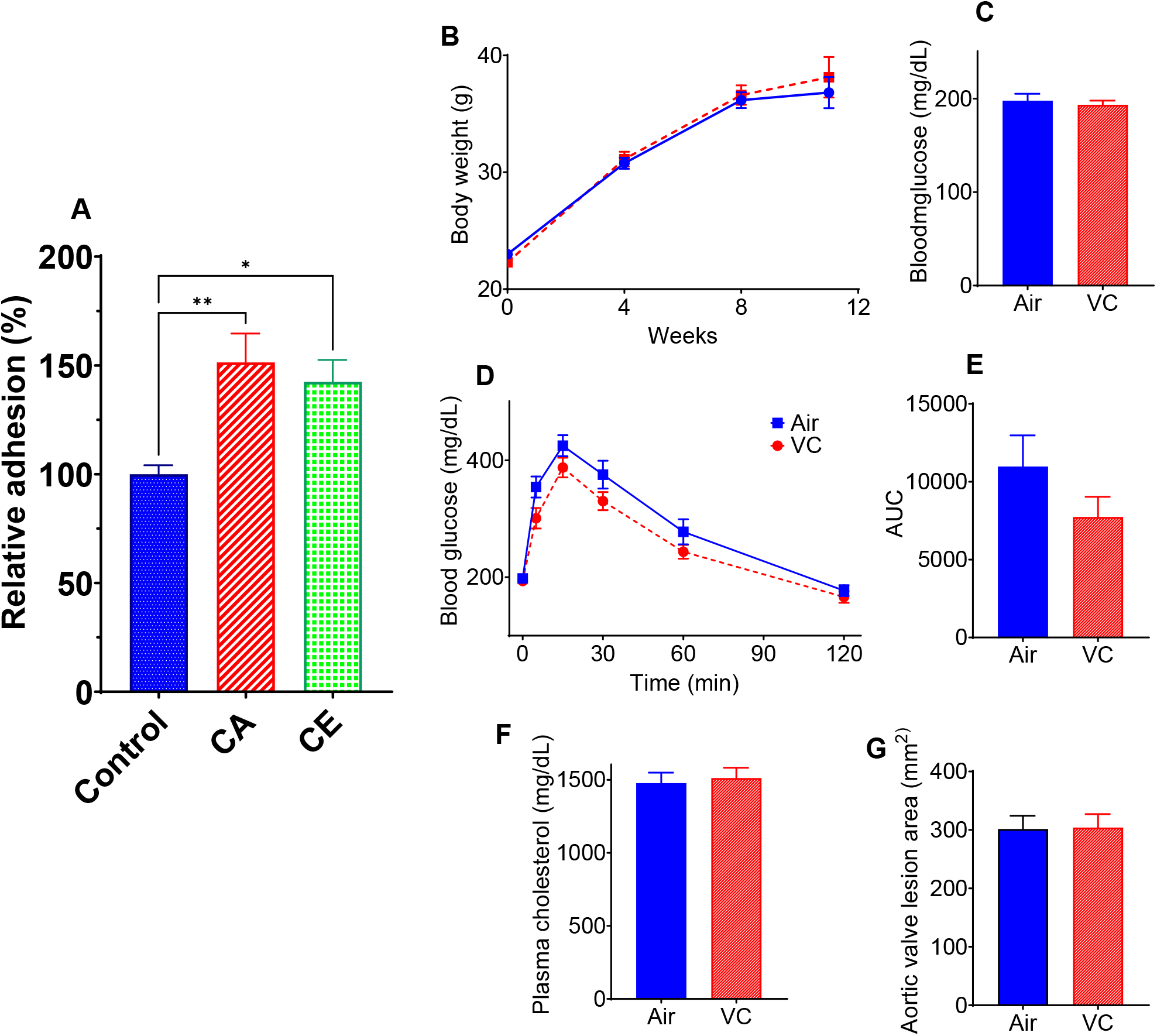
Atherosclerotic lesion formation in vinyl chloride-exposed mice. (**A**) Vinyl chloride metabolites 2-chloroacetaldehyde (CA, 5μM) and 2-chloroethanol (CE, 5μM) increase the adhesion of murine bone marrow derived macrophages to mouse aortic endothelial cells. (**B-G**) Eight-week-old male *Ldlr*-KO mice were maintained on Western diet (WD) for 12 weeks, and body weights (**B**) were measured at indicated time points (*n* = 30/group). (**C**) Blood glucose level at euthanasia. (**D**) GTT in mice exposed to VC for 12 weeks. (**E**) The area under the curve (AUC) for GTT (*n* = 8/group). (**F**) Plasma cholesterol levels. (**G**) Lesions in the aortic valve (lipids were stained with Oil Red O). Values are mean ± SEM. \**P* < 0.05 *vs* air-exposed controls.

## DISCUSSION

Toxicity of high levels of VC exposure has been extensively studied (19–24, 26, 45, 46, 75–78), however, little is known about the effects of chronic low dose VC exposure on cardiometabolic disease. We observed that exposure to 0.8 ppm VC impaired glucose tolerance which was accompanied by the depletion of glutathione and activation of Caspase-1 in the liver, and a modest increase in the infiltration of interstitial macrophages in the lung. Surrogate markers of cardiometabolic toxicity in VC-exposed mice were comparable with controls, and a “second hit” with acute or chronic inflammatory stimuli did not have an additive or synergistic effect on the toxicity of VC. Collectively, these data suggest that although the systemic toxicity of VC is modest, there are localized adverse effects in the liver and the lung.

The primary route of exposure to VC is inhalation, with close to half of inhaled vinyl chloride being absorbed in blood and tissues (79, 80). Absorbed vinyl chloride is metabolized mostly in the liver via oxidation by cytochrome P450 2E1 to form a highly reactive intermediate 2-chloroethylene oxide, which spontaneously rearranges into 2-chloroacetaldehyde (81–84) and then converted to 2-chloroethanol. 2-chloroethylene oxide, 2-chloroacetaldehyde, and 2-chloroethanol conjugate with cellular glutathione to facilitate their excretion. Recent studies suggest that 2-chloroethanol can induce insulin resistance in mice maintained on high fat diet, potentially in oxidative stress- and endoplasmic reticulum stress-dependent manner (46). Our data suggest that chronic exposure to VC significantly decreases hepatic glutathione. Depletion in reduced glutathione levels is sufficient to impair glucose tolerance (85). We recently showed that exposure to benzene induces insulin resistance by depleting hepatic glutathione, augmenting oxidative stress, and potentiating inflammation in mice (64). However, our data suggest that although VC exposure depletes hepatic glutathione, it does not affect the levels of lipid peroxidation products such as MDA. The observed depletion in glutathione levels is likely to be due to the conjugation with the VC metabolites rather than a reflection of oxidative stress-mediated redox signaling or lipid peroxidation. VC exposure did not affect glutathione levels in the lung suggesting that glutathione conjugation of VC metabolites selectively occurs in the liver, where the metabolites are formed. The mechanisms of depletion of HO1 and SOD3 proteins remain unclear. Further studies are required to examine whether VC or its active metabolites induce proteolytic degradation or affect the gene transcription.

The lack of hepatic inflammation in VC-exposed mice suggests that the observed glucose intolerance is not secondary to the inflammatory signaling in the liver of low fat diet-fed mice. Although VC exposure slightly increased the infiltration of interstitial macrophages in the lung, it was insufficient to induce robust pulmonary or systemic inflammation, suggesting that VC-induced glucose intolerance in low fat diet-fed mice is not secondary to pulmonary inflammation. On the contrary, VC induces infiltration of neutrophils, hepatic steatosis, and liver injury in high fat diet-fed mice (46). This may be due to VC-diet interactions.

We have previously shown that environmental pollutants that deplete cellular glutathione and augment oxidative stress (e.g. acrolein and benzene), induce cardiovascular toxicity (86, 87). Exposure to airborne pollutants such as acrolein and benzene is inversely associated with circulating EPC levels in humans (87, 88), and exposure to reagent acrolein and benzene is sufficient to deplete circulating EPCs in mice (86). Acrolein and benzene induce dyslipidemia (9, 86) in mice, acrolein prompts hyper platelet activation (89), and depletes circulating immune cells (49). However, chronic low dose VC exposure in this study did not affect the levels of plasma lipoproteins and circulating EPCs, inflammatory, or pro-thrombotic markers, suggesting that low dose chronic VC exposure does not affect the surrogate markers of cardiovascular toxicity.

To assess whether exposure to VC affects the sensitivity to acute secondary infection, we examined the markers of inflammation in VC-exposed mice. Our previous study showed that VC metabolite 2-chloroethanol potentiates LPS-induced oxidative stress, inflammation, lipid accumulation, and glycogen in the liver (90). However, we did not observe exacerbation of various indices of hepatic, pulmonary, or systemic inflammation in chronic low dose VC-exposed mice treated with LPS. This could be due to the lower dose of LPS used in this study or the chronic exposure to low dose VC rather than a bolus of a specific VC metabolite (90).

To examine the effect of VC exposure on chronic inflammation, we examined atherosclerosis in LDL receptor-KO mice maintained on western diet. Atherosclerosis, the underlying cause of myocardial ischemia and stroke, is the leading cause of death worldwide. It is a chronic inflammatory disease of the arterial wall instigated by the excessive accumulation of lipoproteins and leukocytes in the sub-endothelial space. Repeated failure of innate immune responses to clear sub-intimal LDL, results in the deposition of lipid-laden macrophages or foam cells which secrete pro-inflammatory mediators that facilitate lipoprotein retention and propagate vascular inflammation and atherogenesis (33, 50, 52, 53).

Exposure to airborne industrial chemicals such as acrolein (9) and 1,3-butadiene (10) and an increase in glucose intolerance exacerbates atherosclerosis (33). However, unlike the wild-type C57BL/6 mice, we did not observe compromised glucose homeostasis in the pro-atherogenic LDL receptor-KO mice. Possibly, profound dyslipidemia in the LDL receptor-KO mice conceals the VC-induced relatively modest glucose intolerance. Plasma lipoprotein levels, markers of inflammation and oxidative stress, and atherosclerotic plaque size in the LDL receptor-KO mice in VC-exposed mice were also comparable with air-exposed controls. These observations suggest that unlike other environmental chemicals such as acrolein and arsenic, which induce oxidative stress and vascular inflammation, and exacerbate atherosclerosis (9, 11), low dose chronic exposure to VC does not affect lesional inflammation and atherosclerosis.

Together, our data suggest chronic low-dose VC (at the OSHA-set limit of <1 ppm) induces glucose intolerance, depletes hepatic glutathione, and increases infiltration of interstitial macrophages in the lung of male C57BL/6 mice. However, changes in these biochemical and sub-clinical parameters are insufficient to exacerbate acute or chronic inflammation and atherosclerosis. The OSHA-set limit of <1 ppm exposure appears to have significantly limited the adverse health effects of VC, especially its cardiometabolic toxicity.

## Acknowledgement

This study was supported in parts by NIH grants P42ES023716, 1R01HL149351-01, 1R01HL138992, 1R01HL137229, P20 GM113226, R35ES028373, R01ES032189, T32ES011564, P30ES030283, R21ES031510, 1R01HL149351 and a grant by Jewish Heritage Fund for Excellence OGMN190574L.

## Author Contributions

I.N.Z., B.S.T., T.P.D., W.H.W., I.D.S., B.W., and M.V.M. performed experiments; I.N.Z., W.H.W., M.V.M., M.C.C., and S.S. analyzed data; I.N.Z., M.V.M., and S.S. interpreted results of experiments; I.N.Z., and M.V.M. prepared figures; I.N.Z., and S.S. drafted manuscript; S.S., M.C.C., and I.N.Z. conception and design of research; I.N.Z., B.S.T., T.P.D., W.H.W., I.D.S., B.W., M.V.M, M.C.C., and S.S. approved final version of manuscript.

